# Auxin-dependent alleviation of oxidative stress and growth promotion of *Scenedesmus obliquus* C1S by *Azospirillum brasilense*

**DOI:** 10.1101/2019.12.31.891424

**Authors:** L.A. Pagnussat, G. Maroniche, L. Curatti, C. Creus

**Affiliations:** Laboratorio de Bioquímica Vegetal y Microbiana, Facultad de Ciencias Agrarias, Universidad Nacional de Mar del Plata, Mar del Plata, Buenos Aires, Argentina; Instituto de Investigaciones en Biodiversidad y Biotecnología, Consejo Nacional de Investigaciones Científicas y Técnicas and Fundación para Investigaciones Biológicas Aplicadas, Mar del Plata, Buenos Aires, Argentina; Consejo Nacional de Investigaciones Científicas y Técnicas (CONICET)

**Keywords:** Microalgae, reactive oxygen species (ROS), indole-3 acetic acid (IAA), bacteria, *Azospirillum brasilense*, *Scenedesmus obliquus*

## Abstract

There is currently an increasing interest in the use of microalgae for wastewater treatment and the use of its biomass as a feedstock for biofuels. Both of these applications are often performed more efficiently by microalgal-bacteria consortia. However, the mechanisms that account for the stability and robustness of this kind of interactions are poorly understood. In this study, we confirmed the growth promotion activity of the plant growth-promoting bacterium *Azospirillum brasilense* Sp245 on the microalgae *Scenedesmus obliquus* C1S. We show that this activity is critically dependent on bacterial indole-3 acetic acid (IAA) production, which results in a decrease in algal reactive oxygen species (ROS) levels, higher cell densities and ameliorates algal cells bleaching after nitrogen deprivation. We also show a close inter-species interaction between both partners and an active expression of the bacterial *ipdC* gene involved in production of IAA when co-cultivated.

This study extends the current knowledge of the mechanisms underlying bacteria-microalgae consortia to improve their technological applications and to better understand ecological relationships in the environment.

## 1. Introduction

Microalgae are considered one of the most promising alternative feedstocks to reduce dependency on fossil-based fuels. These organisms accumulate high levels of lipids or carbohydrates that can be used to produce biodiesel or bioethanol, respectively [1, 2]. Additionally, their cultivation has also been considered as a convenient strategy for management of domestic and or/industrial wastewater because of these organisms’ ability to assimilate N, P and organic C from effluents [3, 4]. Under condition of unbalanced growth, especially nitrogen deficiency, most microalgae redirect carbon flow towards accumulation of carbon reserves, mostly triacyclglycerols and/or starch [1, 2, 5, 6]. However, this nutrient limitation causes severe growth reduction which in turn prevents higher biomass, lipids and/or starch productivity [7]. This situation represents one of the most difficult-to-overcome current drawbacks for massive production and commercialization of biofuels from microalgal biomass [8].

Either natural [9] or artificial [3, 9–11] consortia between microalgae and bacteria, have been proposed for both microalgae-based wastewater treatment and or bioenergy production [12]. Microalgal-bacterial consortia frequently outperform pure cultures of either of them, apparently by means of metabolic compartmentalization between the partners enabling pursuit of more complex tasks, which are difficult or even impossible for individual strains or species, and/or improving efficiency of current pathways. Robustness of these consortia is normally enhanced by mutualistic interactions based on need of exchange of specific nutrients and/or growth promoting substances between the organisms involved. It has been proposed that the most common mechanisms by which bacteria exert effects on algal growth are production of vitamins, chelated nutrients and phytohormone-like substances such as IAA, among others [11, 13–17]. More recently, it was shown that the bacterium *Phaeobacter inhibens* promotes growth of the microalga *Emiliania huxleyi* by means of producing low levels of IAA.

However, higher levels of the phytohormone induces algal death by activating pathways related to oxidative stress response and programmed cell death [18].

Algae-bacteria co-culture constitute a promising strategy in biotechnology, as some studies have shown a positive effect of bacteria on algal growth, flocculation, lipid accumulation and cell division [19, 20]. The most thoroughly studied artificial consortium between an algae and a bacterium is perhaps that of *Chlorella* spp. and the plant-growth-promoting rhizobacteria (PGPR) *Azospirillum brasilense,* which produces significant amounts of IAA [21–23]. Experiments with *Chlorella* [19], *Ankistrodesmus* [24] and *Scenedesmus* [25, 26], indicated IAA-dependent responses of growth, chlorophyll content and dry weight. In *Scenedesmus*, IAA supplementation resulted in cell culture synchronization, suggesting an effect on biochemical priming of cellular metabolism that could improve reliability of high density cultivation [27]. In addition, as nutrients become depleted, associated bacteria may help to fulfil the host’s requirements for essential micronutrients, such as chelated micronutrients and cofactors [11, 13, 15].

During the last decade, *Scenedesmus obliquus* arose a one of the most promising oleaginous strains towards biodiesel production [11, 28]. *Scenedesmus obliquus* also engages in a beneficial interaction with *A. brasilense,* which enhances biomass production, coenobium area, colony size, and increases cell density and carbohydrates and proteins accumulation [21, 26, 29].

The aim of this work was to investigate some biochemical aspects underlying the interaction between the oleaginous microalgae *S. obliquus* C1S and the PGPR *A. brasilense* Sp245. We show that bacterial IAA production helps to relieve microalgal oxidative stress under N-limiting conditions.

## 2. Materials and methods

### 2.1 Microorganisms and growth conditions

*Scenedesmus obliquus* C1S (JX014221) was isolated from an artificial pond in the surroundings of Mar del Plata, Buenos Aires, Argentina (38°0’0”S 57°33’0”W) and was used as a study organism . The strain C1S was routinely cultivated in BG11 medium (0.04 g·L^− 1^ K_2_HPO_4_; 0.075 g·L^− 1^ MgSO_4_·7H_2_O; 0.036 g·L^− 1^ CaCl_2_·2H_2_O; 0.006 g·L^− 1^ citric acid; 0.006 g·L^− 1^ ferric ammonium citrate; 0.001 g·L^− 1^ EDTA (disodium salt); 0.02 g·L^− 1^ Na_2_CO3, and trace metal mix (2.86 mg·L^− 1^ H_3_BO_3_; 1.81 mg·L^− 1^ MnCl_2_·4H_2_O; 0.222 mg·L^− 1^ZnSO_4_·7H_2_O; 0.39 mg·L^− 1^ NaMoO_4_·2H_2_O; 0.079 mg·L^− 1^ CuSO_4_·5H_2_O and 0.049 mg·L^− 1^ Co(NO_3_)_2_·6H_2_O), containing 0.5 mM NaNO_3_ (nitrogen-limiting condition) as a nitrogen source. All experiments were inoculated with 5.00 × 10^4^ microalgal cells (counted under light microscope in a Newbauer chamber) and 5.00 × 10^6^ of *A. brasilense* cells (estimated by optical density at 600 nm, OD_600_) in 125 mL flasks with 25 ml of BG11 medium. Cultivation was performed at 30 °C under constant cold white light at an intensity of 100 μmol.m^−2^ .s^−1^ and with manual stirring twice a day. Determinations were performed after 3, 8 and 15 days of co-culture. Three to four replicates were used for each culture condition. For auxin treatments, IAAor 2,4-dichlorophenoxyacetic acid; 2,4-D were applied at final concentrations ranging from 0.0001 to 1 μg.mL^−1^ . Both reagents were purchased from Sigma-Aldrich (USA).

Starter single-species cultures of *A. brasilense* Sp245, the isogenic strain Faj009 [30] (IAAd), Faj009/pRX-PipdC (IAAdPipdC) and Sp245/pOT1e-*pipdC-egfp* [31, 32] were cultivated in Luria-Bertani medium (LB) at 30°C for 18 h with orbital shaking (100 rpm). When required, kanamycin 25 μg mL^−1^(IAAd) or kanamycin plus tetracycline at 25 μg mL^−1^ (IAAdPipdC) were added to the media.

### 2.2 Recombinant strains

*Azospirillum. brasilense* Faj009 - here named IAAd - is a Tn5-insertional mutant of Sp245 in which the *ipdC* gene was interrupted, resulting in a 90% reduction of IAA secretion as compared to the *wild type* strain [30]. The mutation of *A. brasilense* IAAd was complemented using the plasmid pRX-PipdC which contains a functional copy of the *ipdC* gene from *A. brasilense*. To obtain this plasmid, a DNA fragment comprising the gene promoter and the *ipdC* coding region was amplified by PCR using primers Ab.P-*ipdC* F (5’-GGATCCACCCCTCCACAATTTCCGG-3’) and Az39.ipdC R (5’-CTACTCCCGGGGCGCCGC-3’) from the *A. brasilense* Az39 genome. The amplified product was cloned into pGemT Easy (Promega, USA) and sub-cloned into the *EcoRI* site of pRX. To obtain the expression plasmid pRX, pME6000 [33] was digested with *SphI* and *KpnI*, filled-in with Klenow and re-ligated with T4 DNA ligase. The resulting plasmid, devoided of its *SacI* site after digestion and Klenow treatment, was sequentially treated with *EcoRI,* Klenow and *PstI* to introduce the *tac* promoter, which was excised from pME6032b with *EcoRV* and *NsiI*. pME6032b is a derivative of pME6032 [34] in which the *lac* repressor gene was eliminated by treatment with *BamHI*, Klenow and *SspI*, followed by re-ligation. The resulting plasmid pRX-PipdC was introduced into *A. brasilense* IAAd by tri-parental mating using the helper plasmid pRK600 [35] to generate strain IAAdPipdC (Fig 1 A).

**Fig 1.**
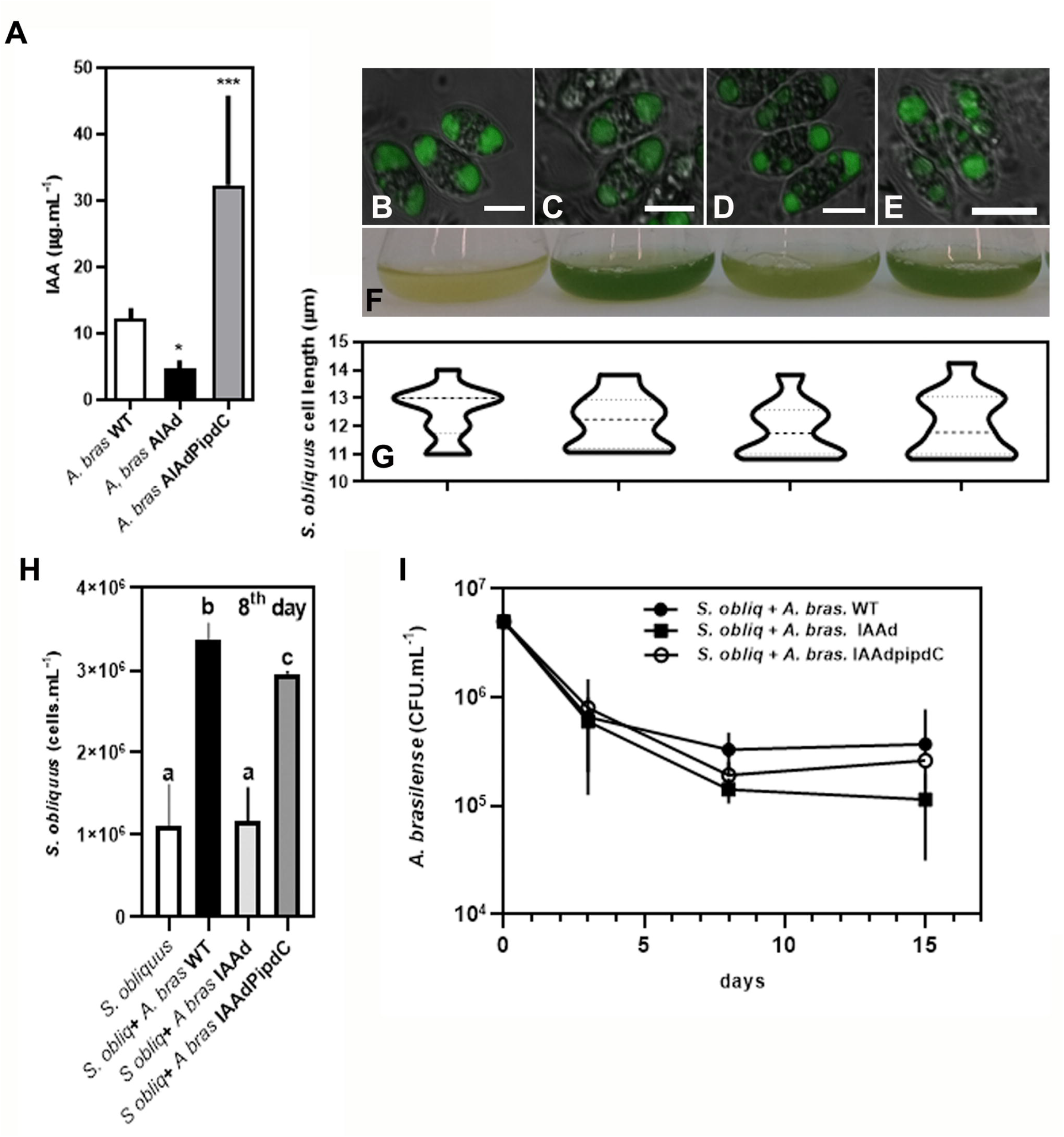
*A. brasilense wt*, IAAd or IAAdPipdC viability and effect on *S. obliquus* C1S under nitrogen limiting conditions. **A**) Comparison of IAA contents in culture media of wt, IAAd and IAAdpiPdC strains of *A. brasilense.* Data represent the mean and SD of three independent experiments ± SD. * represents a significant difference compared to the wt strain (P ≤ 0.05) and ***(P ≤ 0.001). (**B-H**) Representative photographs of **B**) Axenic *S. obliquus.* or *S. obliquus* inoculated with **C**) *A. brasilense wt*, **D**) IAAd or **E**) IAAdPipdC strain. **F**) Aspect of the culture, **G**) cell length (μm) and **H**) density (cells. mL^−1^) of *S. obliquus* after 8 days cultured, from left to right: alone, with *A. brasilense wt*, IAAd or IAAdPipdC. **I**) Bacterial viability on co-culture treatments (CFU. mL^−1^) Plotted data are the means of three independent experiments ± SD. Different letters represent significant differences (P ≤ 0.05).

To generate the *ipdC* reporter strain, plasmid pOT1e-*ppdC-egfp* carrying a *ipdC*::*egfp* transcriptional fusion [31, 32], kindly provided by Dr. Claire Prigent-Combaret from Lyon University, France, was introduced into Sp245 *wild type* strain by tri-parental mating.

### 2.3 Scanning electron microscopy (SEM)

For SEM observation of microalgae and bacteria, samples of three biological replicates for each treatment after three days of co-culture were harvested by centrifugation at 8,000 × *g* for 15 min at 4 °C and the supernatants were removed. Cell fixation was performed for 12 h at 4 °C in the fixative solution containing 0.1 M phosphate-buffered solution (pH=7.3), 2% (v/v) glutaraldehyde, and 4% (v/v) paraformaldehyde and then were washed with deionized water. Samples were then dehydrated with increasing concentrations of ethanol: 50%, 70%, 90% and 100% (v/v) and finally dried in 100% ethanol. The microscopic structure of microalgae-bacteria complexes was observed using a JEOL JSM-6460LV microscope.

### 2.4 Confocal laser scanning microscopy

Microscopic analysis of three biological replicates for of *A. brasilense* Sp245/ pOT1e-*ppdC-egfp* co-cultured for 8 days with *S. obliquus* was performed using a Nikon C1 confocal laser scanning microscope. All images were acquired with a 60x/1.40/0.13 oil-immersion objective. EGFP protein was excited at 488 nm and fluorescence was detected at 515-530 nm. Images were analyzed with Nikon EZ-C1 Freeviewer software.

### 2.5 Nile red staining

Cells were stained with Nile Red (5◻μg.mL^−1^ final concentration; Invitrogen) according to Cakmak (2012) . Images were acquired with a 60x/1.40/0.13 oil-immersion objective, using a Nikon C1 confocal laser scanning microscope. The Nile Red signal excited at 488◻nm and the fluorescent signal was collected between 560 and 600◻nm; chlorophyll fluorescence was captured using a laser excitation line at 633◻nm, and the emission was collected at 515-530 nm. Images were merged with Nikon EZ-C1 Freeviewer software.

### 2.6 ROS determination

Oxidative stress was assayed by measuring dichlorofluorescein (DCF) oxidation according to da Costa *et al*., (2016)[36] Briefly, samples of 3-day-old *S. obliquus* cultures subjected to different treatments were mixed with 4 μM dichlorodihydrofluorescein diacetate (H_2_DCFDA). H_2_DCFDA is de-esterified inside cells to form a free acid that can then be oxidized to the fluorescent DCF. After a 60 min exposure to H_2_DCFDA at 37°C in the dark, fluorescence was measured using a Fluoroskan Ascent fluorescence plate reader (Thermo Labsystems). The excitation filter was set at 485 nm and emission filter at 538 nm. Background fluorescence (no H_2_DCFDA added) was subtracted and the results normalized to 10^6^ microalgal cells. mL^−1^, as determined by flow cytometry. Reported values (means ±SD) correspond to three independent experiments, each one performed with three technical replicates.

### 2.7 Analytical and Statistical Methods

Algal growth was determined by counting the cell number in a flow cytometer Partec CyFlow® space with a blue diode pumped solid-state Laser (20. mW, UNMdP) after 3, 8 and 15 days of cultivation. To determine the size of *S. obliquus* cells, a light microscope (Leica, Germany) and the imageJ software for imaging analysis were used. The size of the cells was studied on the 3^rd^ and 8^th^ day of cultivation. Bacterial colony forming units (CFU. mL^−1^) was determined after 3, 8 and 15 days of co-cultivation with microalgae, by the plate serial dilution spotting method, according to Jett (1997). For IAA quantification, bacteria were grown in 125-ml Erlenmeyer flasks containing LB medium supplemented with 100 μg.mL^−1^ tryptophan until OD_600_ 1, centrifuged and IAA quantified in the supernatant by the Salkowski assay (Mayer 1958) [37] Reported values (means ±SD) correspond to three independent experiments. Results were analysed using Prism 8.02 (GraphPad; Carlsbad, CA) software. Normality and statistical significance of differences between treatments were determined by Shapiro-Wilk and one-way ANOVA with Tukey’s multiple comparisons test using Prism 8.02.

## 3. Results and Discussion

### 3.1 *S. obliquus* C1S growth promotion by *A. brasilense* requires bacterial auxin production

To analyse the effect of *A. brasilense* Sp245 inoculation on *S. obliquus* C1S cultures and more specifically, the involvement of IAA in this interaction, co-cultivation experiments were conducted under nitrogen limiting conditions. Cultivation was performed with *A. brasilense* wild type (*wt*), IAA deficient strain (IAAd) or IAAd complemented strain, which has been transformed with a plasmid containing the *A. brasilense ipdC* gene under its native promoter (IAAdPipdC) (Fig 1 A). Fig 1H shows that after 8 days, *S. obliquus* C1S inoculation with *A. brasilense* Sp245 increased algal growth by 3-fold compared to the control (P < 0.0001). On the other hand, *A. brasilense* IAAd failed to enhance *S. obliquus* growth (Fig 1 H). IAAd revertant strain rescued the failure to promote algal cell growth and prevented algal bleaching after long term incubation under N-limiting conditions (Fig 1 F and H). It should be noted that also under nitrogen sufficiency conditions, co-cultivation with A. brasilense, also promoted microalgal growth (Online Resource 1). As shown in Fig 1, none of the *A. brasilense* strains affected *S. obliquus* cells shape or size significantly (Fig 1B, C, D, E and G). Microalgal longitudinal cell length without bacteria reached 12.1 μm ± 1.14, and under co-culture with *A, brasilense wt*, *A. brasilense* IAAd or *A. brasilense* IAAdPipdC the average size was 11.87± 0.99, 12.17 ± 0.96 and 12.5 ± 0.97, respectively (Fig 1 G). After 15 days under N-limiting conditions, whether they were co-cultered or not with bacteria, microalgal cells exhibit TAGs accumulation in oil bodies, that can be clearly stained with Nile Red (Fig 1 B, C, D and E).

During co-cultivation with microalgae, bacterial viability decreased approximately 10-fold after 3 days, and afterwards they remained unchanged in time. There were no statistically significant differences in bacterial viability among the different bacterial strains (Fig 1 I).

These results indicate that auxins are required for *A. brasilense*–mediated microalgal growth promotion, as has been established for *Chlorella* spp. and other *Scenedemus* strains [21, 22, 26]. In accordance, co-culture of *S. obliquus* with an *A. brasilense* strain expressing *egfp* under the *ipdC* gene promoter, confirmed the active expression of that gene during the assayed culture conditions (Fig 2). Moreover, SEM images also suggested that both the *wt* and the IAAd strains are interacting with the algal surface (Fig 3a and b). This suggests a strong potential for interaction between *A. brasilense* and some microalgae, regardless of its artificial nature. This interaction could result in a more efficient exchange of nutrients and signals (including IAA) between the partners, favouring molecules-mediated interspecific communication. In agreement with these observations, it has been reported that the type VI secretion system (T6SS) of *A. brasilense* is upregulated by IAA [38]. As a consequence, the IAAd strain had a significant reduction in the expression of T6SS components that is reversed by exogenous IAA treatment [38]. Interestingly, the T6SS structural components form an injection tube through which proteins can be injected into a eukaryotic cell and is described as involved in bacterium–eukaryotic cells interaction[38]

**Fig 2.**
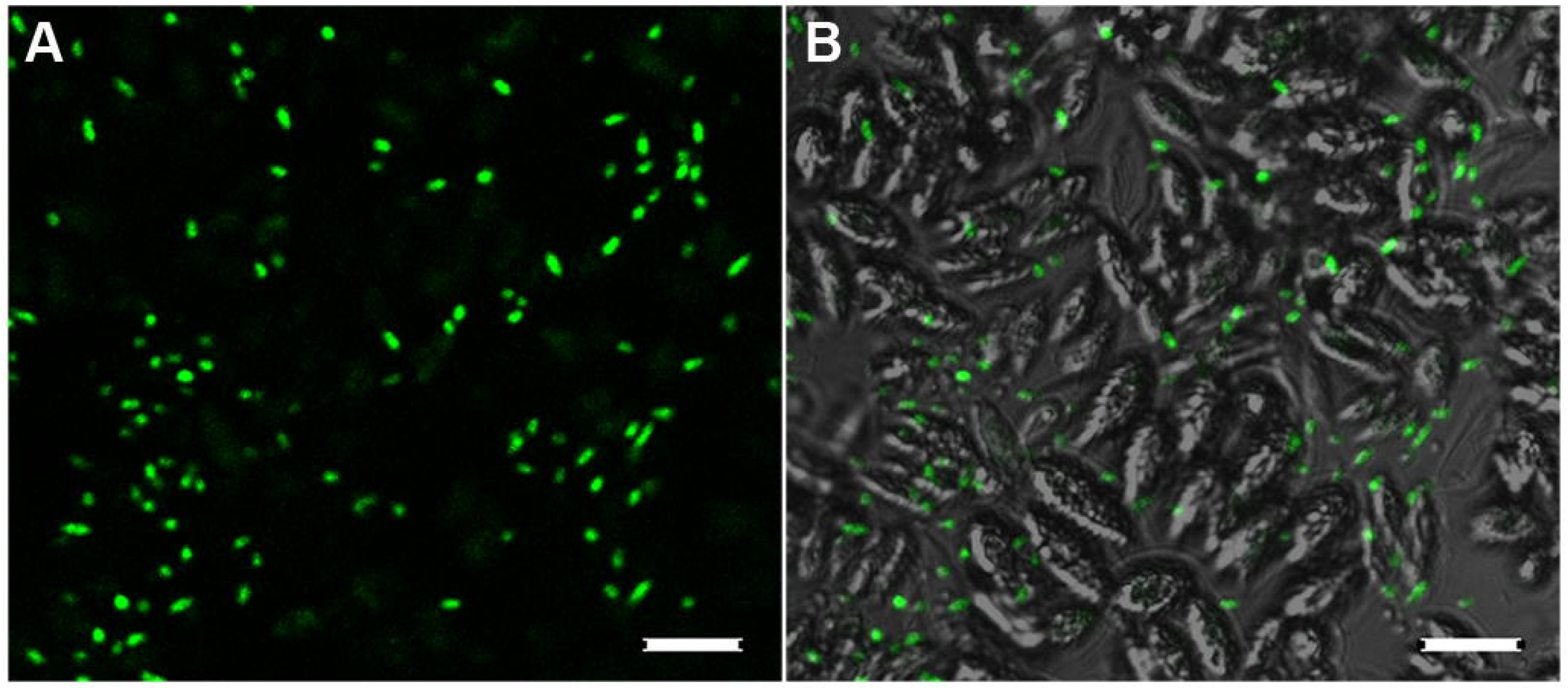
Confocal laser scanning microscopy of *S. obliquus* co-cultured with an *A. brasilense* reporter strain, expressing *gfp* under the *ipdC* gene promoter. **a**) fluorescence image and **b**) merging between fluorescence and bright field images. White bar legth represents10 μm.

**Fig 3.**
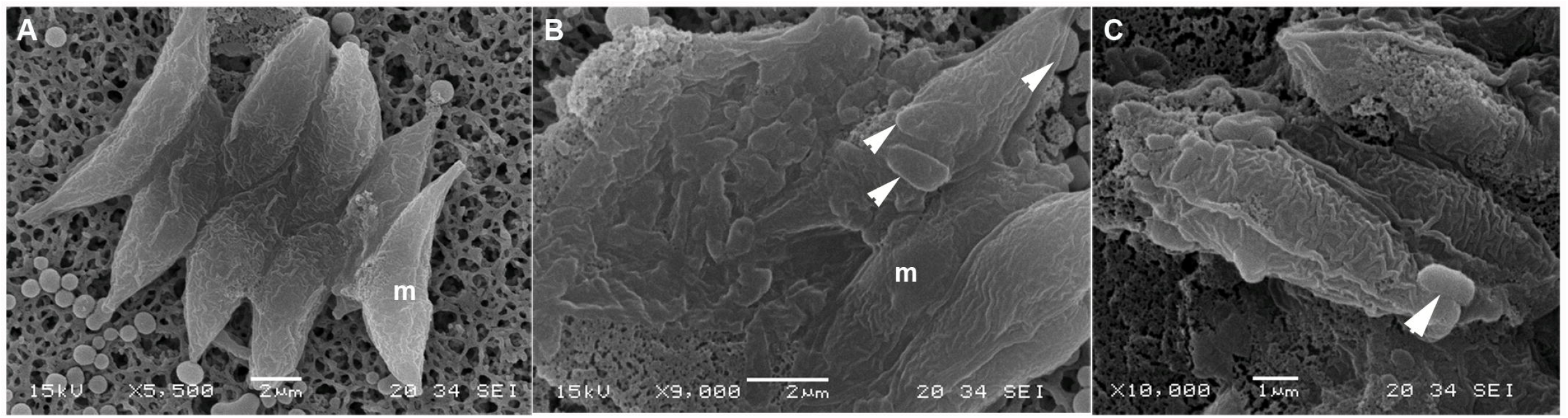
Scanning electron micrography (SEM) of *S. obliquus* cells inoculated with *A. brasilense* strains. **a** S. obliquus cells cultured alone, **b** *S. obliquus* inoculated with *A. brasilense wt* or **c** *A. brasilense* IAAd strain. Representative images were captured after 3 days of co-culture. Arrows point to bacterial cells; m: microalgae. Bar scale: 2 μm.

To further confirm the effect of IAA on *S. obliquus* C1S growth, the exogenous application of the pure chemicals was analysed. Auxins stimulated *S. obliquus* growth only at 0.1 ng . mL^−1^, as revealed by the increase in cell density (Fig 4). The influence of both IAA and 2,4-D on microalgal growth was statistically significant (P < 0.0001 and P< 0.05, respectively) at 3 days of cultivation (Fig. 4a and b). At the experimental endpoint (8 days), 0.1 ng mL^−1^ 2,4-D induced the maximal growth of 1.9 × 10^6^ cells . mL^−1^, a 1.4-fold increase, compared to the control (1.3 ×10^6^ cells . mL^−1^, Fig 4b). Auxin concentrations higher than 0.1 ng . mL^−1^ produced a strong detrimental effect on algal growth (Fig 4).

**Fig 4.**
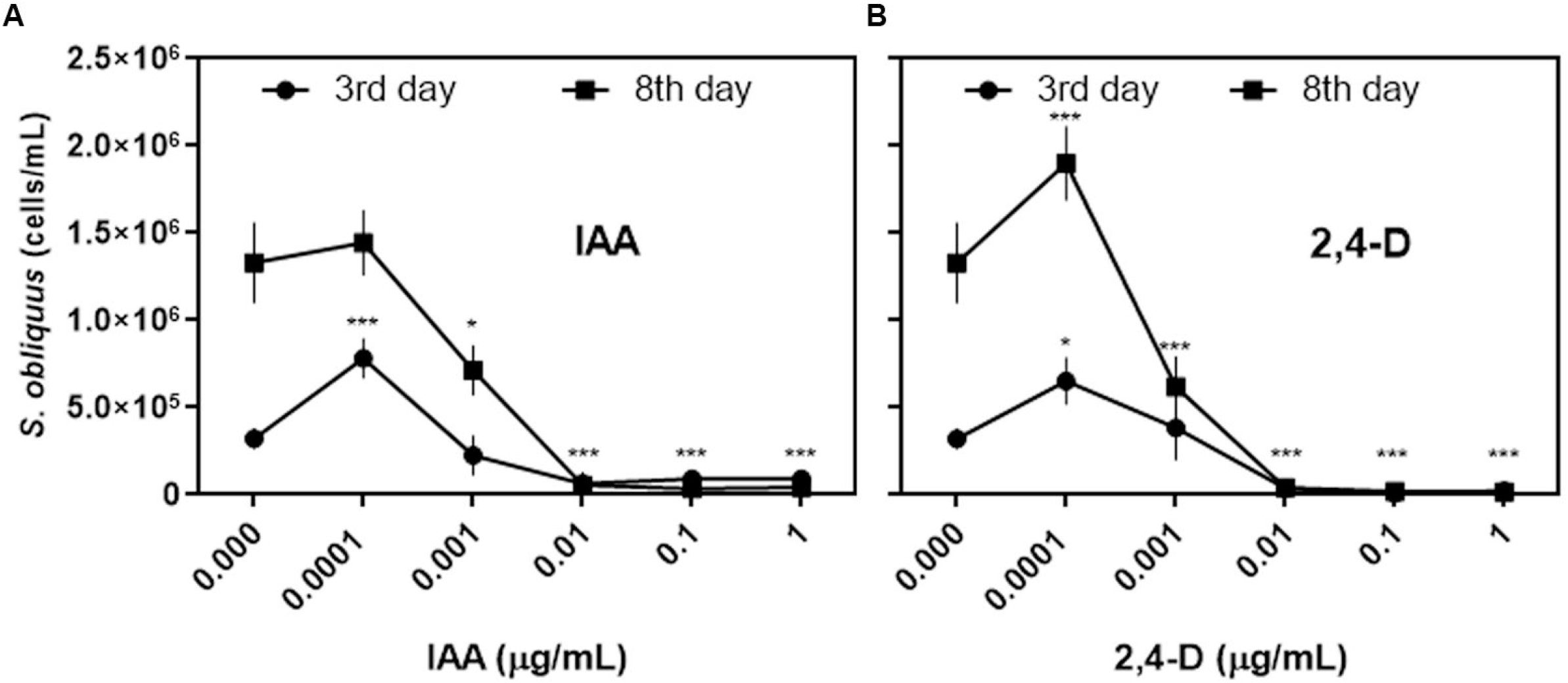
Effect of exogenously applied auxins on *S. obliquus* growth. *S. obliquus* C1S cell density (number of cells/mL) in cultures under the influence of auxins (**a** IAA and **b** 2,4-D) at a range of concentrations of 0 −1 μg/mL after 3 and 8 days of algal cultivation. Data are the means of three independent experiments◻±◻SD. Asterisks represent significant differences * (P ≤ 0.05) and ***(P ≤ 0.001).

Taken together these results confirm that growth promotion of *S. obliquus* C1S by *A. brasilense* Sp245 is mostly mediated by bacterial IAA production.

Previous studies carried out by De-Bashan *et. al*. (2008) and more recently by Choix *et. al*. (2018), showed that the immobilization of *A. brasilense* with *Chlorella sorokiniana* or *S. obliquus* co-immobilized in alginate beads increase microalgae growth and lipid content, as well as carbon fixation in the presence of high concentrations of CO_2_ [21, 22]. Here we showed that co-culture between *A. brasilense* and *S. obliquus* is sufficient to exert microalgal growth promotion, without the need of immobilization in alginate beads, as reported before [21].

### 3.2 *A. brasilense* Sp245 alleviates *S. obliquus* CS1 oxidative stress in an auxin-dependent manner

In plants, the production and detoxification of ROS are regulated by signal transduction pathways that comprise nitric oxide, calcium and auxins, among other signal molecules [10, 39]. The occurrence of these signal transduction pathways in microalgae remains poorly understood. Under natural environmental conditions, algal cells have to cope with ROS generated in mitochondria and chloroplasts, due to respiratory and photosynthetic electron transfer, respectively. While under conditions of nitrogen availability microalgae display a set of coordinated responses to cope with ROS production, when nitrogen is limited, the rate of ROS production to ROS detoxification is altered and cell growth is compromised [13](Online Resource 2b). To further investigate the relationship between ROS production in microalgae and bacterial auxins, we determined intracellular levels of ROS in *S. obliquus* cells co-cultivated with *A. brasilense* or synthetic IAA, upon three different physiological stages 3 days (low growth rate: 4.38 × 10^4^ cells.mL^−1^. day^−1^), 8 days (high growth rate: 2.3 × 10^5^ cells.mL^−1^. day^−1^) and 15 days (oil accumulation, growth rate: 0 cells. mL^−1^. day^−1^). Figure 5 shows that ROS levels, on algal cell basis, remained almost constant up to 15 days of culture. Modification of growth by supplying limiting amounts of N increased ROS levels only marginally at longer incubation times (Online Resource 2). However, inoculation with *A. brasilense* or supplementing exogenous IAA dramatically changed intracellular ROS content in the microalgae. These treatments produced a short-term increase in intracellular ROS at day 3 of cultivation that decreased sharply at longer incubation times, as cell density increased (Fig 5a and b). A strong negative correlation was observed between cell density and ROS production, which might explain the microalgal growth stimulation observed in the presence of *A. brasilense* and synthetic IAA (Online Resource 3).

**Fig 5.**
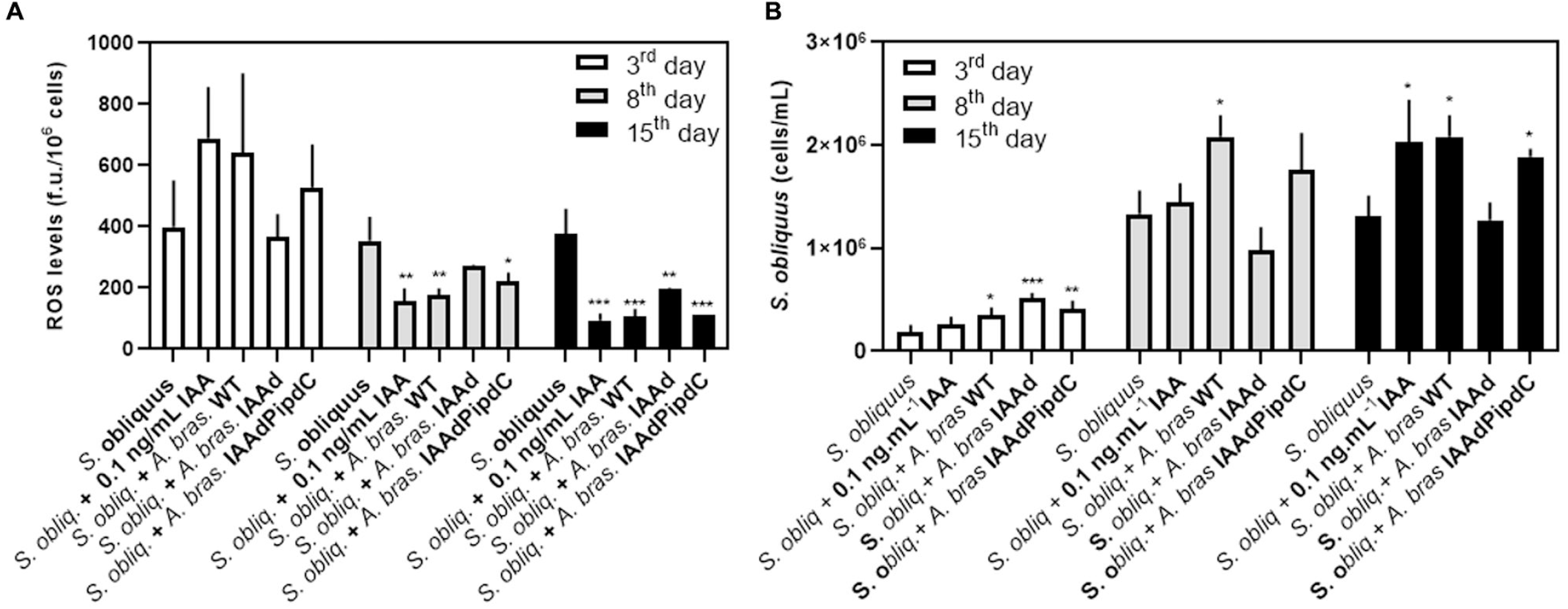
Effect of *A. brasilense* inoculation on ROS production by *S. obliquus* **a** Quantitative analysis of ROS production by fluorometry under different experimental conditions. **b** *S. obliquus* growth on each treatment. Reported values (means ±SD) correspond to three independent experiments, each one performed with three technical replicates. Asterisks represent significant differences * (P ≤ 0.05), ** (P ≤ 0.01) and ***(P ≤ 0.001).

Figure 5a shows a 2-fold decrease in microalgal intracellular ROS after 8 days of co-cultivation with *A. brasilense* Sp245. Exogenous IAA supplementation also diminished ROS levels by 2.25-fold compared to the axenic culture (Fig 5a). On the other hand, co-culture of *S. obliquus* with *A. brasilense* IAAd produced the same ROS levels than the individual *S. obliquus* axenic culture (Fig 5a). As expected, the complemented strain IAAdPipdC behaved similarly to the *wild type* strain and diminished ROS levels by 1.6-fold (Fig 5a). After 15 days, culture with *A. brasilense wt,* IAAdPipdC strains, or exogenous IAA, showed a 3.57, 3.41 and 4.07-fold decrease in microalgal ROS, respectively (Fig 5a). These results indicate that although cell density alone appeared not to alter ROS production by the algae, bacterial IAA increase cell density and decrease ROS cellular levels (Fig 5). Currently, we cannot distinguish whether oxidative stress alleviation enables a higher algal cell density, growth rate promotion by the bacterium results in lower oxidative stress, or both mechanisms are operating concertedly.

Co-culture with the *A. brasilense* strain IAAd, also produced a statistically significant decrease in microalgal ROS (P < 0.001), suggesting that other factors besides IAA may be involved in *A. brasilense*-mediated oxidative stress reduction (Fig 5a). Accordingly, our laboratory previously demonstrated that *A. brasilense* IAAd exerts a weaker plant growth promotion than the *wild type* strain on inoculated tomato plants.

This growth promotion was completely inhibited by a nitric oxide (NO) chelator (cPTIO), suggesting that auxins and NO could act concertedly for plant growth stimulation [40]. In S. obliquus, the addition of a NO donor in combination with H_2_O_2_ prevents the loss of chlorophyll and growth inhibition generated by oxidative stress [41, 42]. In C. vulgaris, NO treatment increased the activity of antioxidant enzymes, lowering ROS levels and providing protection against cell damage caused by herbicides [42]. Since NO exerts its effect at a short distance due to its gaseous nature, physical interaction between the bacteria and microalgae, as the one revealed in the present work (Fig 4), might favour NO signalling in an aquatic environment.

In contrast to the relatively high diversity of microbes observed in the rhizosphere, microalgae surroundings (i.e., phycosphere) are generally colonized by a less complex microbial community. In general, less than 30 bacterial species have been found populating the algal phycosphere [34, 35]. Interestingly, the frequency of co-occurrence between IAA-producing bacteria and green algae in natural and engineered ecosystems has been widely reported [14, 39]. Most of the reported mutualistic interactions involve vitamin-synthesizing bacteria and phytoplankton, nitrogen-fixing cyanobacteria or diatoms and phytoplankton that depend on nearby bacteria for ROS detoxification [43, 44]. Our results are in good agreement with recent studies which showed that the exogenous application of IAA or cytokinins induce the production of antioxidant enzymes (superoxide dismutase and catalase) in *Chlorella sorokiniana* DPK-5 [45] and *S. obliquus* [46] which might diminish ROS levels under nitrogen stress. Our study further extends this biochemical regulation of algal cell metabolism to bacteria-algae interaction.

In sum, our results might not only have some biotechnological implications for uses of bacteria-microalgae consortia for different purposes, but also to help to understand community dynamics in the environment.

## 4. Conclusions

Algae-bacteria consortia have advantages over pure cultures for a diversity of biotechnological applications. The underlying mechanisms for this kind of interactions are already starting to emerge. This study contributes novel pieces of knowledge regarding an IAA-dependent pathway for bacteria-algae molecular signalling resulting in algal growth promotion by alleviation of oxidative stress.

## 5. Statement of Informed Consent

No conflicts, informed consent, or human or animal rights are applicable to this study

