## Supplemental Figures for "Auxin-dependent alleviation of oxidative stress and growth promotion of *Scenedesmus obliquus* C1S by *Azospirillum brasilense*"

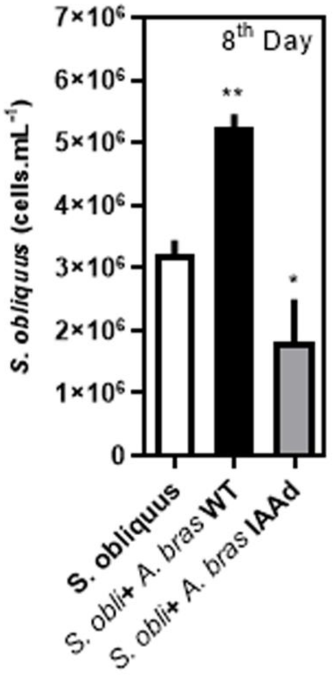


**Online Resource 1** *S. obliquus* cell density upon coculture with *A. brasilense* under nitrogen sufficiency conditions (2 mM Sodium nitrate). Reported values (means ±SD) correspond to three independent experiments, each one performed with three technical replicated. Asterisks represent significant differences * ($P \leq 0.05$), ** ($P \leq 0.01$).

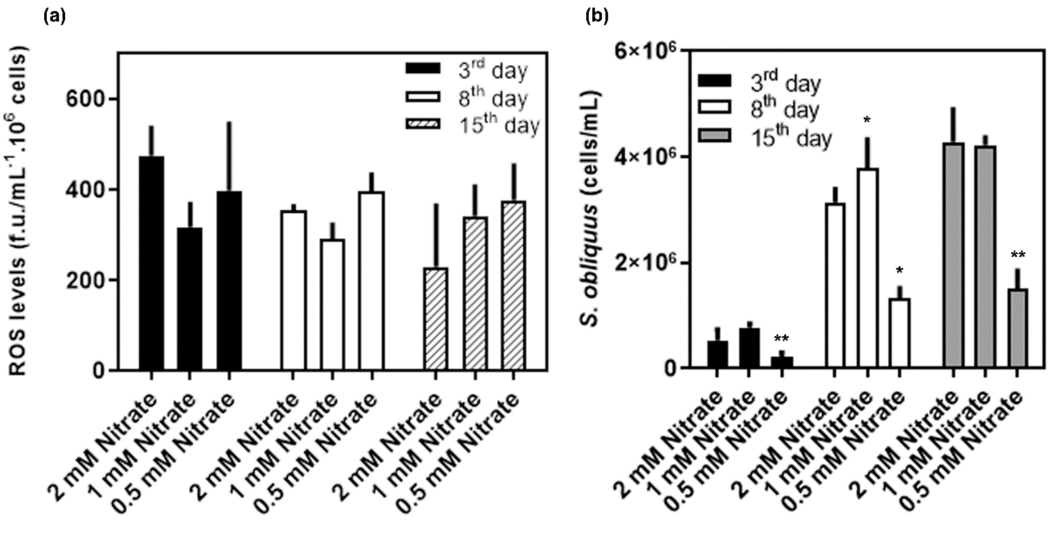


**Online Resource 2** ROS production by *S. obliquus* coculture in three initial nitrate contents a Quantitative analysis of ROS production by fluorometry under different experimental conditions. b *S. obliquus* growth on each treatment. Reported values (means ±SD) correspond to three independent experiments, each one performed with three technical replicated. Asterisks represent significant differences * ($P \leq 0.05$)** ($P \leq 0.01$).

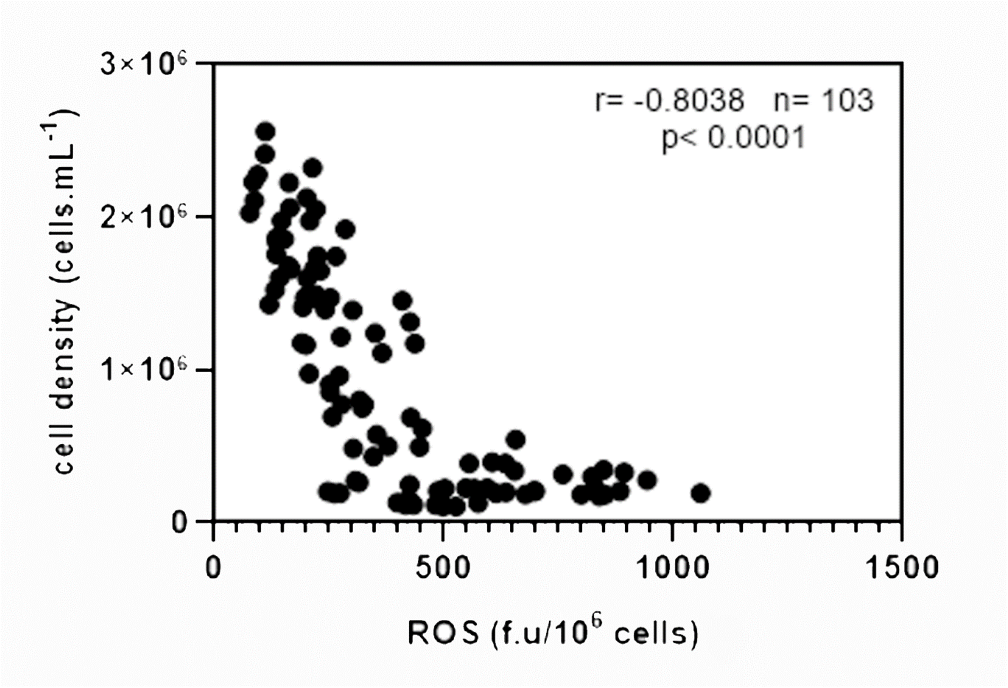


**Online Resource 2** Spearman correlation plot between the microalgal cell density and intracellular ROS levels. Spearman´s coefficient (r), p-value (two-tailed, p) and number of events (n) are shown.
